# Multi-omic dereplication of antibiotic production by diffusion chamber isolated bacteria from Australian soils

**DOI:** 10.1101/2025.07.30.667785

**Authors:** Negero Gemeda Negeri, Duong Duc Anh Nguyen, Miwchael Michael, Emma Kenshole, Calum Walsh, Louise Judd, Liam K. R. Sharkey, Weiguang Zeng, Timothy P. Stinear, Marion Herisse, Chuan Huang, Max J. Cryle, Sacha J. Pidot

## Abstract

Soil bacteria are a major source of clinically useful antibiotics, yet the majority of soil-dwelling microorganisms remain uncultivable by standard laboratory methods. To access this untapped microbial diversity, we employed microbial diffusion chambers to isolate bacteria from ten Australian soil samples. A total of 1,218 bacterial isolates were recovered, representing a diverse collection spanning 61 genera from 32 families, covering the major known phyla of soil bacteria. Antibiotic activity screening revealed that 16% of isolates inhibited the growth of at least one of *E. coli* or *S. aureus*, with 120 isolates displaying activity against multidrug-resistant pathogens including methicillin-resistant *Staphylococcus aureus* (MRSA) and vancomycin-resistant *Enterococcus faecium* (VRE). Mass spectrometry-based dereplication using GNPS identified known antibiotics in 33% of bioactive strains, including actinomycin D, nonactins, and valinomycin. Genomic analysis confirmed the presence of corresponding biosynthetic gene clusters (BGCs), while targeted analysis of selected strains uncovered production of additional antibiotics such as nigericin and streptothricin that were not initially detected by mass spectrometry. Our results demonstrate that diffusion chambers enhance bacterial recovery from soil and show the benefits of a combined pipeline including bioactivity screening, mass spectrometry, and genomics for effective antibiotic dereplication and discovery.

**Impact Statement:** This study delivers a significant advance in natural product discovery by demonstrating that microbial diffusion chambers can dramatically improve the recovery of diverse soil bacteria with antibiotic-producing potential from Australian soil samples. By integrating in situ cultivation with high-throughput screening, mass spectrometry-based dereplication and genome mining, this study yielded more than 1,200 bacterial isolates—including many with activity against multidrug-resistant pathogens—and confirms the production of both known and previously undetected antibiotics. The discovery of streptothricin production via genomics, despite its absence from mass spectrometry data, underscores the power of a multi-layered dereplication strategy. This work not only validates the utility of diffusion chambers for unlocking the "rare biosphere" of uncultivable microbes but also highlights critical methodological refinements to diversify antibiotic-producing strains beyond *Streptomyces*. The findings will inform and accelerate future efforts to discover urgently needed antibiotics from environmental microbiomes.

**Data Summary:** The whole-genome sequences were deposited in the National Center for Biotechnology Information National Library of Medicine under BioProject accession number PRJNA1272227.

## Introduction

Soil microbiomes represent an immense reservoir of yet-to-be-discovered bacterial diversity, with the untapped potential to yield novel antibiotics and bioactive compounds. Indeed, soil bacteria have been the most prominent source of antibiotic molecules, with the majority of current clinically used antibiotics derived from these organisms (1). With the rise in antimicrobial resistant pathogens comes an urgent need to continue finding new antibiotics. While soil has been a great source, it is estimated that less than 1% of soil bacteria can be cultivated under standard laboratory conditions, severely limiting our ability to access this rich diversity of microorganisms and creating a significant barrier to uncovering new species and discovering novel bioactive compounds (2, 3).

To address this issue, microbial diffusion chambers have emerged as an effective tool for in situ cultivation, enabling the isolation of previously uncultivable bacteria by mimicking their native environment (4, 5). These chambers permit the exchange of nutrients and signalling molecules with the surrounding soil microbiome while containing the target microorganisms for subsequent cultivation and analysis. Previous studies have demonstrated that this approach can significantly increase the diversity of cultivable bacteria from environmental samples and potentially access previously untapped reservoirs of secondary metabolites (6, 7).

Once isolated, bacteria must be systematically investigated for antibiotic activity. Classical cell-based assays provide an excellent method to prioritise antibiotic producing microbes that are active against any chosen organism. However, identification of bioactivity alone is not sufficient to determine the identity of the antibiotic that is produced, and this requires efficient methods to distinguish novel antibiotics from the extensive catalogue of known antimicrobials. Advances in mass spectrometry-based dereplication coupled with molecular networking tools such as Global Natural Products Social Molecular Networking (GNPS) have revolutionized the dereplication process, allowing researchers to rapidly identify known compounds and prioritize novel chemical entities for further investigation by comparing spectral data against known molecular databases (8, 9). This approach aids in distinguishing novel compounds from previously identified antibiotics, streamlining the discovery process. Additionally, the integration of genomic approaches to identify genes for antibiotic production, known as biosynthetic gene clusters (BGCs), provides crucial information about the potential of isolated strains to produce novel antimicrobial compounds and can confirm the results of mass-spectrometry investigations (10).

In this study, we employed microbial diffusion chambers to isolate bacteria from Australian soil samples and evaluated their potential to produce antibiotic compounds. We utilized a comprehensive workflow combining bioactivity screening, mass spectrometry-based dereplication, and genomic analysis of biosynthetic gene clusters to identify and characterize antibiotic-producing isolates. Our results demonstrate the efficacy of this integrated approach for the discovery of both known and potentially novel antimicrobial compounds, while providing insights into the environmental factors that regulate their production.

## Methods

### Strains, media and culture conditions

All recovered isolates were cryopreserved in 20% glycerol at -80°C. Reasoner’s 2A (R2A) agar or broth, and SMS agar were made according to Berdy *et al* (6). Soft nutrient agar was made with a final agar concentration of 0.6% agar for antibiotic overlay assays. A panel of antibiotic sensitive (*Escherichia coli* DH10B and *Staphylococcus aureus* Newman) and multi-drug resistant bacteria (*Escherichia coli, Klebsiella pneumoniae, Staphylococcus aureus, Enterococcus faecium,* and *Acinetobacter baumannii*) were used as test organisms (Table S1).

### Collection of soil samples

Soil was collected from ten sites in Victoria, Australia at a depth of approximately zero-15cm from the soil surface. Topsoil and any accompanying foliage was collected, transported and stored in a clean plastic bag. For incubation of prepared diffusion chambers, the soil sample was collected up to 24 hrs before use and stored at room temperature until use.

### Construction of bacterial diffusion chambers

Bacterial diffusion chambers were prepared using a silicone sealant (acetic cure), 0.03 µm semipermeable membranes (Whatman Nucleopore polycarbonate track-etched membranes, Cytiva) and reused 96-well pipette tip inserts, according to the method outlined in Berdy *et al* (6). Membranes were cut to a size approximately 2cm larger than the surface area of the 96-well pipette tip insert; the overhanging membrane was sealed to the side of the pipette insert, ensuring a thorough seal around the diffusion chamber. Pipette tip inserts were sterilised by submersion in 70% ethanol and allowed to dry before bottom membranes were attached. Diffusion chambers with a bottom membrane affixed were irradiated with UV light for 20 minutes in a laminar flow hood and were then left inside the laminar flow hood for up to 16 hours until fully cured. Following inoculation (see below), the top of the membrane was sealed with another piece of 0.03 µm semipermeable membrane. Diffusion chambers were then left in the laminar flow hood for up to 1 hour for the top membrane sealant to dry, before being placed in their soil sample for incubation.

### Inoculation and cultivation of diffusion chambers

To inoculate diffusion chambers, soil slurries were initially prepared as outlined in Berdy et al (6). Cell counts from the microbial fraction of the soil slurries were performed by staining cells with a 1:1000 dilution of Syto9 nucleic acid dye (Thermo Scientific) and 10 µL of each dilution was placed onto a haemocytometer, fixed by evaporation at 37°C for five minutes, coated with 5µL of ProLong^TM^ Gold Antifade Mountant with DAPI (Thermo Scientific) and viewed using a fluorescence microscope (BX53, Olympus).

Molten SMS agar was then inoculated with bacteria at an approximate concentration of one cell per 100µL (based on the estimate from cell counting). Nystatin (Gibco^TM^) was added to inoculated SMS agar at a concentration of 100 U/mL to prevent fungal growth and 100 µL of this SMS agar/bacterial cell mixture was aliquoted into 92 wells of the prepared diffusion chamber. Four wells of the diffusion chamber were inoculated with sterile SMS agar and nystatin (at concentration outlined above) as negative controls. The agar was allowed to set for 30 mins and diffusion chambers were then sealed as outlined above.

Once prepared, each diffusion chamber was placed inside a clean sealed container containing approximately 5cm of soil from the respective soil sample. Soil was moistened with distilled water to allow efficient nutrient transfer (6) and containers were sealed with plastic wrap and incubated in the dark at room temperature for either two, three or four weeks. Following incubation, diffusion chambers were removed from their soil samples in a laminar flow hood and excess external soil was removed by irrigation with sterile water.

### Comparison of diffusion chamber bacterial recovery

At the same time as the diffusion chambers were inoculated and using the same molten agar/cell mixture, wells of a 24-well plate were inoculated with 500µL of the same SMS agar/bacteria/nystatin mixture as outlined above. 24-well plates were maintained in a humid environment at room temperature for the same incubation duration as the diffusion chambers.

### Retrieval and domestication of diffusion chamber colonies

Top membranes were removed from cleaned diffusion chambers, ensuring no debris contaminated individual wells. Agar plugs in each diffusion chamber well were removed, broken up and spread onto sterile R2A agar plates. Plates were incubated at room temperature for up to four weeks and were checked regularly during this time for bacterial growth. On average, each well contained one to three colonies. Individual colonies were picked and passaged onto new R2A agar plates to isolate pure cultures before being stored in cryostorage.

### Antibiotic overlay assays

Isolates were grown from glycerol stocks in 10mL of R2A broth at 28°C with aeration at 200rpm for seven days. From each culture, cells were inoculated into designated wells of a 96-well plate in 100µL aliquots containing 15% glycerol and stored at -80°C until use. 96-well plates were made in triplicate and upon use, were defrosted and used only once before being discarded. A 96 solid pin multi-blot replicator (VP407FP12, V&P Scientific) was used to transfer approximately 1.5 µL of defrosted culture from the wells of each prepared 96-well plate onto duplicate 10×10 cm petri dishes containing R2A agar and incubated at 28°C for up to 14 days. Prior to overlaying, plates were UV irradiated for 20 minutes.

Strains comprising the antibiotic activity test panel are listed in Table S1. Each isolate was grown in 10mL of LB broth (*E. coli* strains) or BHI broth (*S. aureus, K. pneumoniae*, *A. baumannii* and *E. faecium* strains) overnight at 37°C with aeration at 200rpm. The optical density at 600 nm (OD_600_) of each culture was measured and cultures were diluted to OD_600_ 0.6 in 100mL of molten 0.6% nutrient agar. The molten agar-bacteria mixture was poured directly over the UV irradiated test plates (∼40mL per 10×10 cm plate). Overlays were incubated overnight at 37°C. After overnight incubation, plates were visually assessed and photographed for zones of growth inhibition.

Isolates that were positive for antimicrobial activity from the initial overlay screen, were screened a second time individually, using the same protocol as above, to confirm activity. Only isolates showing antimicrobial activity in both initial and secondary screening were deemed positive for antimicrobial activity.

### Pooled 16S rRNA extraction and sequencing

Cells from all 1218 isolates were pooled and nucleic acids were extracted using the Qiagen DNeasy kit according to the manufacturer’s protocol and DNA quality and concentration was assessed using a Nanodrop spectrophotometer and Qubit fluorometer (Thermo Scientific). Full length (V1 - V9) 16S amplicons were generated using the Oxford Nanopore 16S barcoding kit 24 v14 (Oxford Nanopore) according to the manufacturer’s instructions. Sequencing libraries were loaded onto R10.4.1 flow cells. Reads were filtered to only retain those longer than 1400bp and shorter than 1700bp to remove any non-16S sequences. Species-level taxonomy was assigned to the resulting reads by Emu (11).

### Metabolite extraction from diffusion chamber isolates

For dereplication of antibacterial isolates by mass spectrometry, isolates were grown on their producing media for 14 days at 28°C and a small (∼1cm diameter) agar plug was removed from close to the centre of the colony and transferred to a 1.5mL Eppendorf tube containing 600 µL of ethyl acetate. Plugs were sonicated in a water bath for 10 minutes, centrifuged at 13,000 x *g* for five minutes and the supernatant was transferred into a clean 1.5mL Eppendorf tube. Supernatants were dried down in the CentriVap^®^ Benchtop Vacuum Concentrator (Labconco) for 15 minutes at 40°C then resuspended in 400µL of methanol. Resuspended samples were then filtered through 1mL syringes with 0.2 PTFE syringe filters (Thermo Scientific) into clean 1.5mL Eppendorf tubes. Filtrates were kept at -20°C prior to mass spectrometry.

### LC-MS/MS analysis of extracted metabolites

Samples were analysed using LC-ESI-HRMS/MS on a Fusion Lumos Tribrid mass spectrometer (Thermo Fisher). A 2 µL injection volume of samples was introduced by a Dionex Ultimate 3000 UHPLC (Thermo Scientific) fitted with a Kinetex 1.7 µm EVO C18 column (1 mm x 100 mm) (Phenomenex). Compounds were separated using a gradient of water/0.1% formic acid (A) and acetonitrile/0.1% formic acid (B) (2% B to 100% B over 15 min) at a flow rate was 0.075 mL/min. Acquisition was performed in positive ion mode using data dependent acquisition and automatic switching between full scan MS and MS/MS acquisitions. Full scan spectra (m/z 50 – 1500) were acquired in the time of flight (TOF) and the top ten most intense ions in a given scan were fragmented by collision induced dissociation (CID) utilizing stepping.

Thermo .raw data files were converted to .mzXML format using MSConvert, as part of the Proteowizard package (12) and uploaded to the GNPS server (gnps.ucsd.edu) (8). A metadata file (.txt) comprised of sample attributes was also generated and uploaded to GNPS. Molecular networks were generated by GNPS with the following parameters: precursor ion mass tolerance (0.05 Daltons) and fragment ion mass tolerance (0.05 Daltons) and cosine score of at least 0.7. The resulting network was processed further using the Dereplicator (13) and Dereplicator+ (9) algorithms, as implemented in the GNPS environment, to identify known natural products. Dereplicated spectral data were visualised using Cytoscape v3.9.1 (14).

### DNA extraction and whole genome sequencing

Selected strains were cultured in 10 mL sterile ISP2 broth with aeration at 180 rpm and 30°C for at least 7 days. Cultures were centrifuged at 8000 rpm for 5 minutes to collect cell pellets. DNA was extracted using the magnetic bead-based GenFindv3 DNA extraction kit (Beckman Coulter). DNA was prepared for short-read sequencing using the NexteraXT sample preparation kit (Illumina) and was sequenced on a NextSeq500 (Illumina) using 2 x 150 bp reads. DNA was prepared for long read sequencing on the MinION platform (Oxford Nanopore) using the Ligation Sequencing kit. Samples were barcoded using the Rapid Barcoding Kit 96 v14 (Oxford Nanopore).

### Assembly, annotation and bioinformatic analysis of sequenced bacterial genomes

Genomes were assembled using flye 2.9-b1768 (for nanopore reads) (15) or the assembly script shovill (https://github.com/tseemann/shovill), which uses the SPAdes v3.15.0 assembler (16) (for Illumina reads). All assemblies were annotated with Prokka (17). For species identification, the 16s rRNA sequences were extracted from annotated whole genome assemblies and were searched against the rRNA/ITS database at NCBI using BLASTn. The hit with the highest percentage identity was selected as the best match. Secondary metabolite biosynthetic gene clusters (BGCs) were predicted using AntiSMASH v7 (https://antismash.secondarymetabolites.org) (10).

### Nigericin analysis

Extracts from strain 10362 were prepared by growing bacteria on R2A agar plates for 14 days, cutting the agar into pieces, transferring to a clean flask and extracting with ethyl acetate (100ml). The contents were mixed and incubated at room temperature overnight. The ethyl acetate fraction was then poured over anhydrous sodium sulfate and filtered with Whatman #1 filter paper (GE healthcare) into a round bottom flask. Solvent was evaporated under vacuum using a rotary evaporator (Buchi) with a water bath set at 40°C. Dried extracts were resuspended in 1ml HPLC grade methanol and stored at 4°C.

The LC chromatographic analysis was performed on 1290 Infinity III (Agilent) equipped with a reverse-phase C18 column (Kinetex 2.6 µm C18 100 Å, 75 x 3 mm, Phenomenex). The column temperature was maintained at 40°C, with a flow rate of 0.5ml/min. Standard nigericin and bacterial extracts were separated using a 20min gradient composed of 100% MS grade water (solvent A) and 100% MS grade acetonitrile (solvent B), both containing 0.1% formic acid. The gradient was as follows: 0-8min, 20% B; 8-12 min, 100% B; 12–15 min, 100% B; 15–16 min, 20% B; 16–20 min, 20% B. The LC-eluent was introduced into the mass spectrometer (Revident Q-TOF (Agilent)) with a dual ESI as an ionisation source, operating in positive-ion mode. Data acquisition and processing were performed using Agilent MassHunter software. A commercial nigericin standard (BioAustralis) was used to confirm the retention time and exact mass of nigericin and as a comparator for putative nigericin containing extracts. Ion chromatograms were extracted for *m/z* 742.5150 [M+NH^4^]^+^ (calculated for C_40_H_72_NO_11_ = 742.51054).

### Determination of streptothricin production

Antibiotic plug-based assays were performed by growing strain 10258 on R2A agar plates for 14 days at 30°C. Small plugs of mycelial growth (∼0.5cm diameter) were cored out and were placed on top of an LB agar plate that had been inoculated with either *E. coli* BW25113, *E. coli* BW25113 (pGDP1:*stat)* (streptothricin resistant) or *E. coli* BW25113 (pGDP1:*uvrA*) (echinomycin resistant) (18). Any zones of clearing were observed and recorded.

## Results

### Isolation and cultivation of bacteria from microbial diffusion chambers

Based on previous reports suggesting that microbial diffusion chambers could improve bacterial recovery and provide access to novel taxa compared to standard plating techniques (3, 5), we attempted to construct diffusion chambers to investigate bacteria present in Australian soil samples. Several types of diffusion chamber devices have been developed (3, 5, 19, 20) and we based ours upon the method of Berdy et al (6), using easy to obtain laboratory materials including semi-permeable membranes and inserts from standard tip boxes. Following construction of the diffusion chambers, we tested whether they could increase bacterial recovery compared to standard cultivation methods. We inoculated the wells of diffusion chamber devices and miniaturised petri dishes with the same soil bacteria/agar mixture that had been diluted to approximately 1 cell/100 µL of agar. Following growth for at least 3 weeks, we then counted the number of colonies present in the diffusion chamber and the petri dishes and converted these to cells recovered per 1 mL of agar. Diffusion chambers provided superior bacterial recovery compared to plate-based cultivation, increasing colony counts by at least 3-fold (Fig. 1A).

**Fig. 1.**
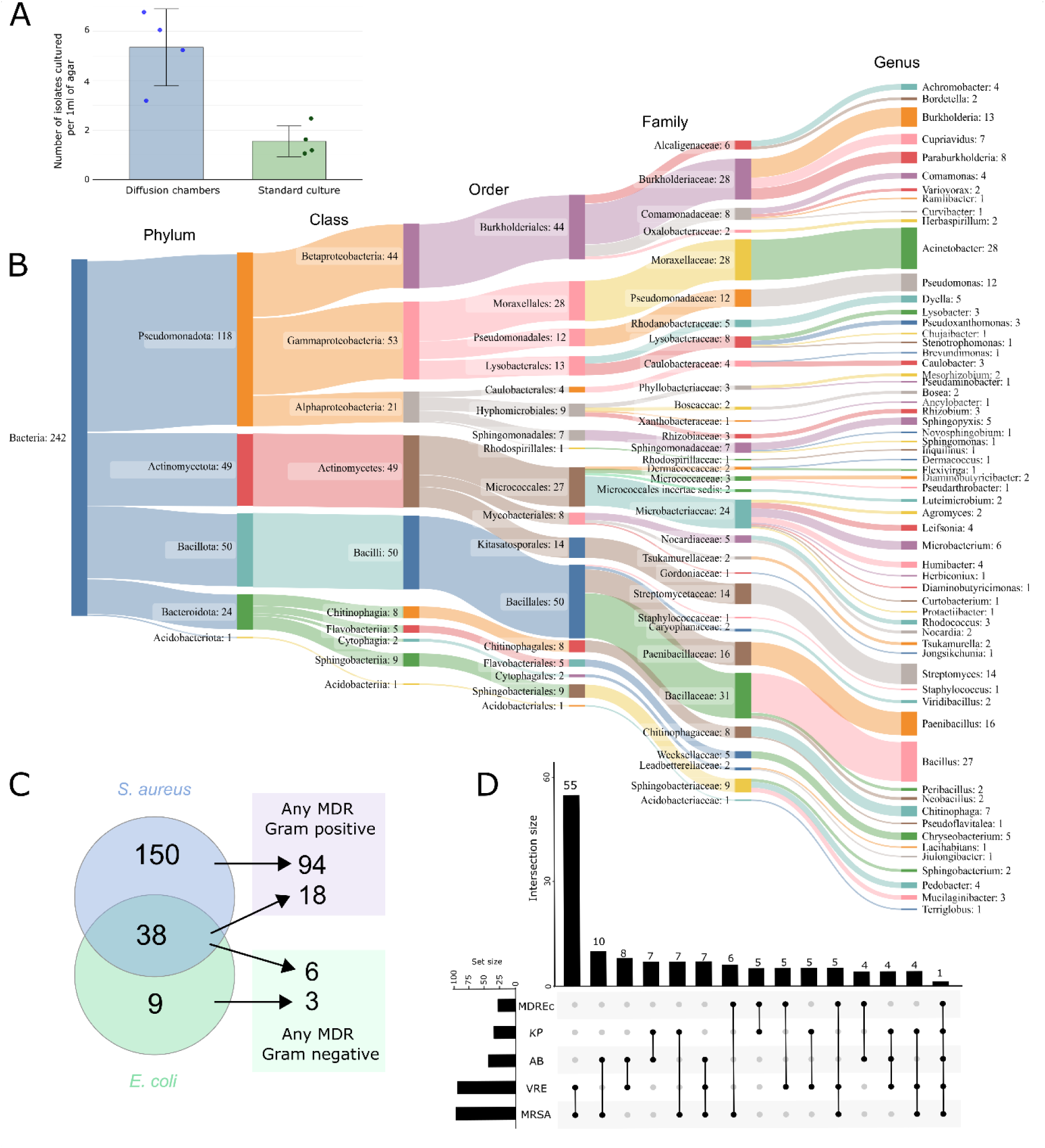
Cultivation and bioactivity data for diffusion chamber isolated bacteria. A) Comparison of diffusion chambers to standard cultivation method shows increase in bacterial recovery using diffusion chambers. B) Sankey plot showing diversity of isolated bacteria, as estimated from bulk, full-length 16S rRNA sequencing. C) Venn diagram showing numbers of isolated bacteria exhibiting antibiotic activity against antibiotic-sensitive *S. aureus* Newman or *E. coli* DH10B. Arrows point to number of strains from each subset that were active against one or more Gram positive or Gram negative MDR pathogen. D) Upset plot showing number of isolates that exhibited bioactivity against one or more multidrug resistant pathogens. MRSA, methicillin resistant *Staphylococcus aureus;* VRE, vancomycin resistant *Enterococcus faecium*; AB, *Acinetobacter baumannii*; KP, *Klebsiella pneumoniae*; MDREc, multidrug-resistant *E. coli*.

Based on these results, we used diffusion chambers to attempt to recover bacteria from Australian soil samples. From 10 different soil samples, we isolated a total of 1218 bacterial isolates using diffusion chambers. Following incubation in their respective soil samples, bacterial microcolonies were visible within the wells of the diffusion chambers within 14 days and the isolated organisms exhibited a wide variety of morphologies upon purity plating. To rapidly gain an insight into the diversity of the isolated bacteria, we performed bulk long read 16S rRNA sequencing on all isolates. Amplification, sequencing and subsequent deconvolution of the 16S rRNA gene sequences through the Emu pipeline (11) revealed 242 non-redundant sequences from five bacterial phyla, corresponding to 10 different classes and 32 different families (Fig. 1B). At the genus level, 61 different genera were identified, including genera that are commonly found in soil, such as *Streptomyces* and *Bacillus*. Although our bulk sequencing strategy did not reveal the identity of every isolated organism, approximately 20% of our cultivated isolates were identified, which was sufficient to show that diverse organisms from a range of phyla and families can be isolated using diffusion chambers.

### Antibiotic activity of diffusion chamber-derived bacteria

Given that most of our current antibiotics have been identified from soil bacteria, we sought to determine the extent of antimicrobial production in our isolated bacteria. We tested our collection for antibiotic activity against an antibiotic-sensitive preliminary Gram-positive and Gram-negative test set (*S. aureus* and *E. coli*, respectively, Table S1). Of the 1218 isolated bacteria, 197 were found to have any antibacterial activity (16%), with 38 isolates showing broad spectrum activity, 150 showing only anti-*S. aureus* activity and nine showing only anti-*E. coli* activity (Fig. 1C). Isolates displaying antibacterial activity against drug sensitive strains were then tested for their ability to inhibit antibiotic resistant pathogens, including methicillin resistant *S. aureus*, *Klebsiella pneumoniae*, *Acinetobacter baumanii*, *E. coli* and vancomycin-resistant *Enterococcus faecium*. Of the 197 isolates exhibiting activity against either of the antibiotic-sensitive organisms, 115 isolates (58% of antibiotic-producing bacteria) inhibited at least one multidrug resistant pathogen. The majority of these isolates had activity against MRSA and/or VRE, while inhibitors of any of the Gram-negative MDR pathogens were relatively rare (Fig. 1D). Of those with broad spectrum activity in our initial screen, only one was capable of inhibiting all test strains, while the majority of strains with anti-*S. aureus* activity in the initial screen were capable of inhibiting either MRSA and/or VRE (94 of 150, 63%). For those strains with only initial anti-*E. coli* activity, three isolates were identified that could inhibit any one or more of the multidrug resistant Gram-negative pathogens.

### Dereplication of antibiotic producing isolates

With the large number of antibiotic producing bacteria that were isolated, we sought to rapidly identify any known antibiotics produced by these organisms. To this end, we attempted dereplication by collection of HR-ESI-LC-MS/MS spectra for each isolate exhibiting antibiotic activity and comparing these data against known spectra. Initially, we collected mass spectral data for 192 isolates and compared these spectra to those from known molecules using the Global Natural Products Social Molecular Networking (GNPS) tool (8, 21). GNPS generates a molecular network where nodes represent ions detected by a mass spectrometer and edges connect molecules with similar MS/MS fragmentation spectra, suggestive of similar molecular structures. Within the GNPS suite of tools, Dereplicator and Dereplicator+ (9, 13) directly compare the uploaded spectra to a curated set of reference spectra. Doing so, allowed the identification of a presumptive known natural product in 65 isolates (33% of the tested strains).

The most commonly identified antibiotic was actinomycin D (also known as dactinomycin), which was found by GNPS in extracts from 12 of the 65 dereplicated isolates. The majority of actinomycin D producing isolates were also found to produce the related congener actinomycin X2 (also known as actinomycin V). Both molecules are known to have antibiotic activity against Gram positive bacteria, which correlates well with the results of the bioactivity assays for their producing strains (Table S2). Eleven strains were found to produce antibiotics from the nonactin group of compounds, including bonactin, nonactin, dinactin and trinactin, all of which are also known to have antibiotic activity against Gram positive bacteria (22). Valinomycin was another commonly identified antibiotic (produced by ten isolates) and was found to be produced by four strains that also produced nonactin compounds. Other less commonly identified antibiotics included pyrromycin (produced by six isolates), echinomycin (one isolate), and novobiocin (one isolate). Although we did not test specifically for activity against eukaryotic cells in this study, GNPS identified several known antifungal and antitumour compounds in the extracts of 30 strains, including manumycin (14 isolates), antimycin (nine isolates), montanastatin (seven isolates), tambromycins (five isolates) and venturicidin (one isolate). This analysis shows that we were able to isolate bacteria producing a wide range of bioactive molecules and to use mass spectrometry to dereplicate molecules from 65 isolates (Fig. 2).

**Fig. 2.**
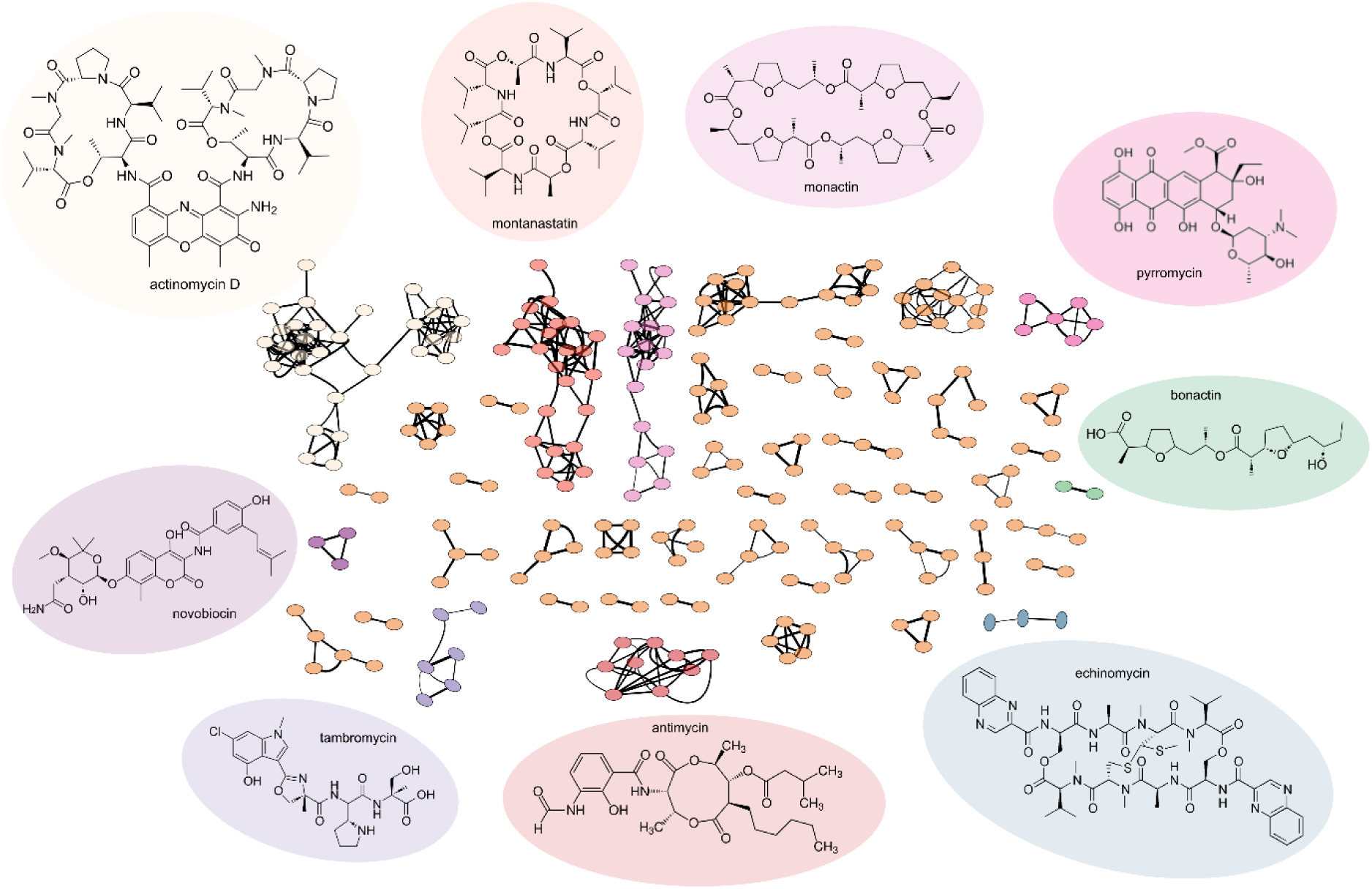
Selected dereplicated compounds detected in ethyl acetate extracts of diffusion chamber derived isolates. The GNPS derived molecular network shows known compounds and their derivatives as molecular families (all nodes joined together within a subnetwork represent a molecular family). Known molecules are shown and their background is colour coded according to the subnetwork seen in the diagram.

### Genomic analysis of selected producing isolates for known BGCs

Although mass spectrometry-based dereplication was able to putatively identify a wide range of antibiotic molecules, using this type of dereplication in isolation may result in false positive identifications. Correlation of the biosynthesis genes responsible for the production of known molecules provides confidence that mass spectrometry-based identifications are real. In addition, for strains where antibiotics were not detected by GNPS, analysis of genomes may reveal genes for the biosynthesis of antibiotics that were undetectable by mass spectrometry or not extracted by our choice of solvent.

To investigate the presence of biosynthetic genes for the actinomycin family of molecules, we sequenced the genomes of several producers (Table S3), as the BGC responsible for these compounds is well characterised and should be readily identifiable from genome assemblies. Following genome sequencing and annotation of four actinomycin producing isolates (*Streptomyces murinus* 10260, *Streptomyces antibioticus* M126, *Streptomyces murinus* 10202 and *Streptomyces murinus* 10256, Table S3), assembled genomes were analysed using AntiSMASH v7 (10), a bioinformatic tool to identify secondary metabolite BGCs directly from genome level data. This analysis identified the actinomycin BGC in all four genomes. To determine the extent of actinomycin BGC conservation among our isolates, a comparative analysis was performed with the reference actinomycin BGC from MiBIG (23) (accession BGC0000296.5). This analysis showed that the majority of genes were highly conserved among all actinomycin BGCs (Fig. 3). However, the BGCs in strains 10202 and 10256 contained the insertion of an additional four to six genes directly downstream of the *acmM* orthologue, which encodes a cytochrome P450 monooxygenase (Table S4). In addition, the reference actinomycin BGC from *Streptomyces anulatus* (previously *Streptomyces chrysomallus*) contains an inverted duplication of nine genes at either end of the BGC (24), which was suggested to be an essential part of actinomycin production. However, this inverted duplication is not seen in any of the actinomycin BGCs identified in this study, suggesting that these duplicated genes are not necessary for actinomycin production (Fig. 3A). Furthermore, even though strains 10260, 10202 and 10256 were all identified as *S. murinus* by 16S rRNA analysis, the actinomycin BGCs contained in each strain differ in terms of gene content and organisation. In addition, we were unable identify any significant difference in the actinomycin BGC from producers identified as producing actinomycin D or X or both. These results provide a genetic basis for actinomycin production in isolates 10260, M126, 10202 and 10256, and corroborate the mass spectrometry-based dereplication of these isolates as actinomycin producers.

**Fig. 3.**
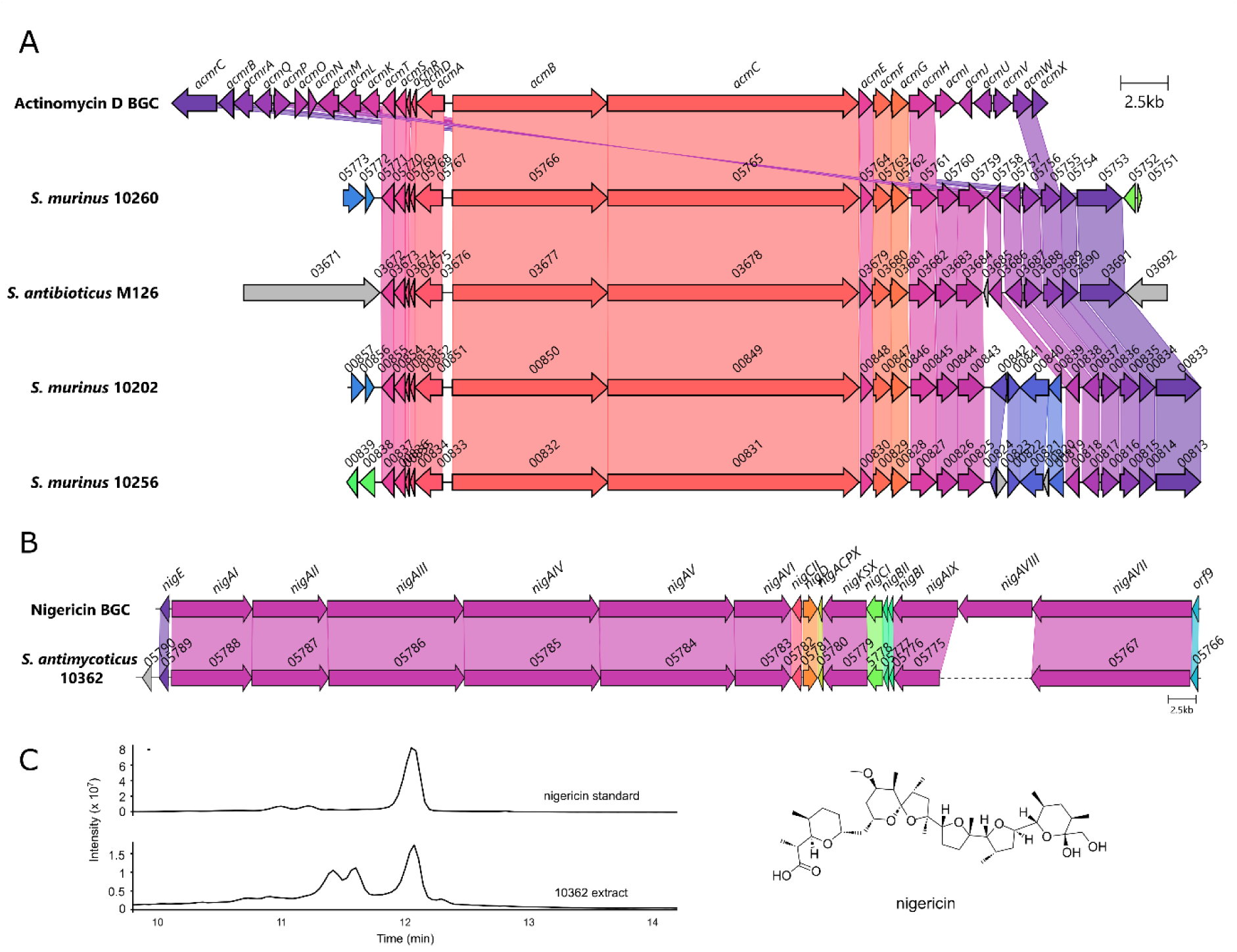
A) Comparative analysis of actinomycin BGCs identified in this study compared to the reference actinomycin BGC. Gene names or locus tags are shown above each gene. Note: *acmrB* to *acmK* are paralogues of *acmH* to *acmX*. Hence, although *acmIJUV* appear to be missing from the diffuson chamber derived isolates, homology can be seen to the paralogous genes *acmLOPQ*. Note also the insertion of at least 4 genes in 10202 and 10256 genomes (blue). B) Homology between the reference nigericin BGC (MiBIG accession BGC0000115) (25) and nigericin BGC from 10362. Gene names or locus tags are shown above each gene. The dotted line in the 10362 BGC indicates a contig break. C) Extracted ion chromatogram from both nigericin standard and culture extract of strain 10362 at *m/z* 742.5150 [M+NH_4_]^+^ and figure showing molecular structure of nigericin.

Confirmation of MS results by genomic data was also possible for strain 10362 (*Streptomyces antimycoticus*), which had broad spectrum antibiotic activity and was found to produce the antibiotics elaiophylin, geldanamycin and BD-12 (Table S2). To investigate the presence of BGCs for these known antibiotics, we sequenced and assembled the *S. antimycoticus* 10362 genome, yielding a large 11.8 Mb genome in 32 contigs (Table S3). AntiSMASH analysis revealed the presence of the complete BGCs for elaiophylin and BD-12 within the 10362 genome, with all genes highly conserved to the corresponding MiBIG reference BGCs (Fig. S1 and S2). For geldanamycin, the BGC was fragmented across at least three contigs in the 10362 assembly, most likely due to high internal sequence homology within the polyketide synthase encoding genes leading to incomplete assembly (Fig. S3). However, all tailoring enzyme encoding genes were highly conserved and syntenous with the reference geldanamycin BGC (Fig. S3). Further investigation of the AntiSMASH report revealed the presence of a BGC for the production of the antibiotic nigericin (Fig. 3B). Although parts of the BGC were located on several contigs, there was sufficient homology to be a clear match to the known nigericin BGC. Nigericin is known to be highly non-polar and it therefore may have been missed by the chromatography conditions used in our rapid extraction and dereplication pipeline. To investigate whether 10362 produced nigericin and whether it too may have contributed to the observed antibiotic activity, we performed targeted extraction and LC-MS analysis, comparing our results to a commercial nigericin standard. Our analysis showed that this strain did indeed produce nigericin on R2A media and that this antibiotic may also have contributed to the observed bioactivity of strain 10362 (Fig. 3C).

### Genomic analysis of isolates producing unidentified antibiotics

We then set out to determine if genome sequencing could help to dereplicate strains where antibiotics were not detected by mass spectrometry. We investigated the genome of strain 10258, which had broad spectrum antibiotic activity, but where no compounds were identified by mass spectrometry-based dereplication. Genome sequencing, assembly and annotation yielded an 8.6Mb genome in four contigs and 16s rRNA analysis assigned this strain as *Streptomyces ardesiacus* (Table S3). Analysis by AntiSMASH v7 revealed the presence of 29 BGCs. While several of the BGCs had 100% homology to well characterised Streptomycete BGCs, including coelichelin and desferrioxamine, the most likely candidate for an antibiotic producing BGC was the streptothricin BGC. Streptothricin has broad spectrum Gram positive and Gram negative activity and is a common antibiotic produced by soil Streptomycetes (18, 26). Comparison of the putative streptothricin BGC in strain 10258 to the reference streptothricin BGC from MiBIG (accession BGC0000432) showed the two BGCs to be identical in gene content and synteny (Fig. 4A).

**Fig. 4.**
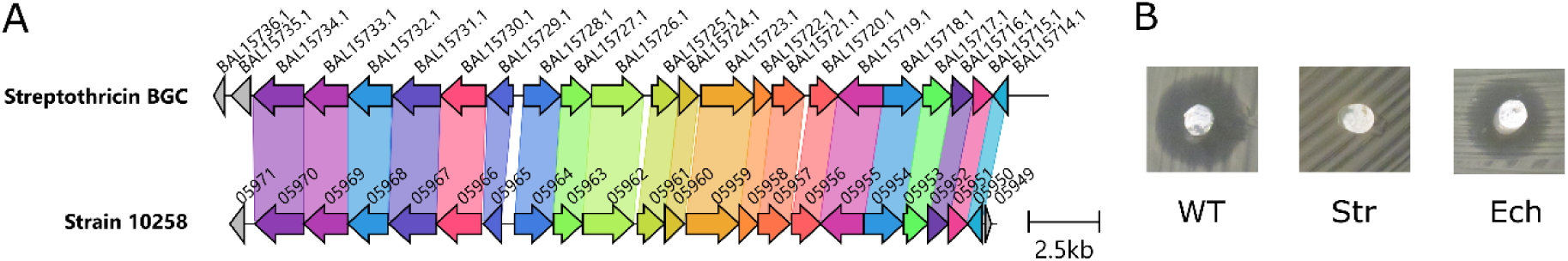
A) Comparative analysis of streptothricin BGC identified in strain 10258 to the reference streptothricin BGC (MiBIG accession BGC0000432). B) Agar plug assay for bioactivity of strain 10258, showing activity against wild-type *E. coli* (WT) and *E. coli* containing an echinomycin resistance gene (Ech), but no activity against *E. coli* carrying the *stat* gene, encoding streptothricin resistance (Str).

While the antibiotic spectrum of streptothricin matches well with the observed bioactivity of strain 10258, we sought to confirm the production of streptothricin through antibacterial assays against an *E. coli* strain containing a streptothricin resistance gene (*stat*), wild-type *E. coli* and *E. coli* bearing resistance to an unrelated antibiotic, echinomycin (18). These experiments showed that strain 10258 formed a distinct zone of clearing against the WT *E. coli* and the echinomycin resistant *E. coli* strains but was incapable of killing the *E. coli* strain harbouring the *stat* gene (Fig. 4B). These results show that strain 10258 contains an intact streptothricin gene locus and that streptothricin is responsible for the antibiotic activity of this strain. These results also show the benefit of combining genomic and targeted microbiological analyses to dereplicate antibiotic producing isolates, as streptothricin was not detected by mass spectrometry alone.

## Discussion

In this study we have utilised microbial diffusion chambers to isolate a large collection of bacteria from Australian soil samples. In total, we succeeded in isolating 1,218 bacterial strains, providing a sufficient number of organisms for our goal of antimicrobial testing. We found that diffusion chambers improved bacterial recovery from soil samples compared to standard plating techniques (Fig. 1A), which others have also noted (3–5, 27). In addition, we showed that our isolated bacteria were diverse, with 242 non-redundant species identified across 61 genera (Fig. 1B). Previous 16S amplicon-based studies of Australian soil samples have found that Proteobacteria, Actinobacteria and Acidobacteria were the dominant members of the bacterial microbiota, with Firmicutes and Bacteroides being present at relatively low abundance (28–31). Our bulk sequencing approach showed that members of all of these bacterial groups were present in our collection (Fig. 1B), although we were unable to estimate recovery percentages for each phylum. Nevertheless, our cultivation approach complements previous sequencing-based studies and to the best of our knowledge is the first report of an isolation-based study of Australian soil microbes with identification to the species level. While the previously mentioned studies all noted Acidobacteria in high abundance in Australian soil samples, we only saw limited evidence of Acidobacterial cultivation; a known problem as these microbes are considered to be very difficult to isolate (32). Future studies aimed at enhancing Acidobacterial cultivation from the environment will be beneficial in further understanding this group of bacteria.

While our pooled 16S rRNA sequencing approach did not allow for the identification of individual strains, it did allow us to assess bacterial diversity of a portion of our isolates, as a surrogate for total diversity within the collection, in a highly time and cost effective manner. Given our bulk DNA extraction approach, it is likely that some samples may have been missed due to limited number of cells within the pool or difficulty in bacterial lysis and DNA extraction from all bacteria using a single method. It has been previously noted that DNA extraction methods can influence the observed diversity in 16S rRNA sequencing studies (33). Therefore, it is possible that the diversity of our isolated microbes is greater than we currently appreciate and further work to individually identify each of the isolated organisms is ongoing.

Notably, 16% of all our isolated bacteria exhibited antibiotic activity, with a significant proportion of these (115 total isolates, 58% of all antibiotic producing bacteria) capable of inhibiting at least one organism in our multidrug-resistant test panel. Inhibition of MRSA and VRE were the most common phenotypes seen, with only a limited number of isolates capable of inhibiting Gram negative bacteria. This lack of anti-Gram negative activity is a known issue in antibacterial discovery projects and is a direct result of molecular challenges in overcoming the outer membrane permeability barrier (34, 35). To rapidly dereplicate these strains, GNPS molecular networking was used, identifying production of a known antibiotic in 33% of tested strains. Producers of actinomycin D and its congeners were particularly prevalent, along with organisms that produced compounds from the nonactin group, valinomycin, and pyrromycin. While our approach rapidly dereplicated many isolates, mass spectrometry-based methods are also susceptible to both false positives (incorrect compound identification) and false negatives (known compounds produced by an organism but not identified because they are not in the mass spectral database). In our case, an additional factor for rapid dereplication is choice of extraction solvent. While many known molecules were identified, it is also likely we missed several due to extraction only with ethyl acetate. As an example, streptothricin appears to have been missed via our ethyl acetate extraction approach. This may be a reason why we only identified one streptothricin producer, while it is estimated that between 1 in 10 to 1 in 1000 of all isolated *Streptomyces* species produce streptothricin (36). Although we were able to use a *stat* (streptothricin acyltransferase) containing *E. coli* strain to test for streptothricin production, previous studies have shown that *stat* does not provide resistance to the related molecule BD-12 (37). This suggests that, while using resistant bacteria to test for the production of known antibiotics may provide a further rapid dereplication step, it must either be combined with further analyses or use a much larger panel of resistant isolates to reduce the chance of spending unwanted time and effort on producers of known antibiotics (18). This reflects the complexity of natural product discovery, where multiple approaches should ideally be combined to enhance identification of truly new molecules.

Focusing our whole genome sequencing efforts on a select panel of antibiotic producing isolates revealed several interesting phenomena. For example, *S. murinus* isolates have been previously shown to produce molecules in the actinomycin complex, as well as lankacidin and lankamycin (38–40). However, none of the three individual *S. murinus* isolates cultivated here possessed the BGCs for either lankacidin or lankamycin. In addition, each isolate was shown to contain the actinomycin BGC in a different genomic context (Fig. 3A). Previous studies have also shown that isolates designated as the same species by 16S rRNA analysis contain different BGCs, and therefore have the potential to produce strain-specific suites of natural products (41). Our work here further confirms these previous observations and suggests that there is still value in evaluating individual isolates regardless of their species designation. In addition, our *S. antimycoticus* isolate was shown to produce both BD-12 and nigericin, which has not previously been described for this species.

As mentioned above, our goal in using diffusion chambers for bacterial isolation was primarily to increase the diversity of isolated soil bacteria, and by doing so, unveil new antibiotic producing microbes. It is sobering, therefore, that all of our sequenced antibiotic producing isolates were identified as members of the genus *Streptomyces*, the best known and most commonly isolated bacterial antibiotic producers (26). While we aimed for an unbiased approach in our colony selection following the domestication of bacteria, it is possible that we selected more *Streptomyces* isolates as these were the most easily identifiable or that they simply grew faster and outcompeted other bacteria. Our experience here suggests that to truly diversify bacterial isolation, diffusion chambers may need to be combined with domestication on a variety of media types (instead of just R2A) to maximise isolation and selection of non-*Streptomyces* bacteria. Cultivation of rare genera from diffusion chambers has previously led to the isolation of novel antibiotic molecules, such as teixobactin and lassomycin (7, 42) and attempts to further diversify bacterial recovery may lead to the discovery of further new molecules. An alternative explanation is that Streptomycetes really are the “best” bacterial antibiotic producers and hence why they are more commonly identified. More recent genomic studies, however, argue against this and have shown that many other bacteria, in particular other members of the phylum Actinomycetales, possess vast untapped secondary metabolic potential (43–46). In either case, future efforts at soil screening should be devoted towards isolating as wide a variety of bacteria as possible.

## Conclusion

Here, we have shown the utility of microbial diffusion chambers for recovering diverse soil bacteria with antibiotic-producing potential. Our integrated workflow—spanning cultivation, bioactivity screening, dereplication, and genomic analysis allowed the identification of strains producing known bioactive compounds. While many antibiotic-producing isolates belonged to the genus *Streptomyces*, our findings underscore the importance of methodical dereplication and strain prioritization in natural product discovery and provide evidence of the need to optimise isolation protocols to diversify bacterial recovery. Furthermore, the detection of bioactivity not captured by mass spectrometry alone reinforces the need for complementary genomic and functional assays. To maximize the potential of diffusion chamber approaches, future efforts should focus on diversifying cultivation conditions to access rarer taxa and novel metabolites.

## Acknowledgements

We thank the staff from the Mass Spectrometry and Proteomics Facility, Bio21 Institute, University of Melbourne for assistance with mass spectrometry data collection.

