## Supplemental figure and tables for "Multi-omic dereplication of antibiotic production by diffusion chamber isolated bacteria from Australian soils"

### Contents

**Table S1.** Strains used in this study
**Table S2**. Bioactivity and compounds Identified by mass spectrometry from the 65 dereplicated isolates.
**Table S3**. Statistics of sequenced bacterial genomes.
**Table S4**. Homology between proteins encoded in actinomycin BGCs.
**Fig. S1**. Homology between the reference BD-12 BGC and BD-12 BGC from strain 10362.
**Fig. S2**. Homology between the reference elaiophylin BGC and elaiophylin BGC from strain 10362.
**Fig. S3**. Homology between the reference geldanamycin BGC nigericin BGC from strain 10362.
**References**

**Table S1.** Strains used in this study

| Strain | Features | Source/reference |
|---|---|---|
| *Escherichia coli* DH10B | F⁻ *endA1 deoR⁺ recA1 galE15 galK16 nupG rpsL Δ(lac)X74* φ80*lacZΔM15 araD139 Δ(ara,leu)7697 mcrA* Δ*(mrr-hsdRMS-mcrBC)* $Str^R$ λ⁻ | Invitrogen |
| *Staphylococcus aureus* Newman | Antibiotic susceptible *S. aureus* strain. | (1) |
| *Staphylococcus aureus* JKD6008 | MRSA, vancomycin-intermediate resistance, aminoglycoside and trimethoprim resistant | (2) |
| *Klebsiella pneumoniae* NTUH-K2044 | Hypervirulent *K. pneumoniae*. Resistant to ampicillin. | (3) |
| *Acinetobacter baumannii* BAA-1710 | Resistant to cefazolin, cefepime, cefotaxime, ceftazidime, ceftriaxone, ciprofloxacin, gentamicin, piperacillin, piperacillin-tazobactam, tetracycline, ticarcillin, trimethoprim-sulfamethoxazole | ATCC |
| *Enterococcus faecium* AUS0233 | Resistant to vancomycin, trimethoprim, tetracycline, macrolides, aminoglycosides | (4) |
| *E. coli* BPH0530 | Resistant to extended spectrum beta-lactams (ESBL), aminoglycosides, tetracycline, trimethoprim, quinolones | (5) |
| *E. coli* BW25113 | *E. coli* K12 derivative. F⁻ DE(*araD-araB*)567 *lacZ4787*(del)::*rrnB*-3 *LAM⁻ rph*-1 DE(*rhaD-rhaB*)*568 hsdR514* | (6) |
| *E. coli* BW25113 (pGDP1:*stat)* | Streptothricin resistant | (7) |
| *E. coli* BW25113 (pGDP1:*uvrA*) | Echinomycin resistant | (7) |

**Table S2**. Bioactivity and compounds Identified by mass spectrometry from the 65 dereplicated isolates

| Strain # | Compounds identified by GNPS, Dereplicator and Dereplicator+ | Observed antibiotic activity (SaN, EcD, MRSA, VRE, Kp, Ab, EcB)* |
|---|---|---|
| 10260 | Actinomycin D | SaN, EcD, MRSA |
| 10202 | Actinomycin D, Actinomycin X | SaN, EcD, MRSA, VRE |
| 10207 | Actinomycin D, Actinomycin X | SaN, EcD, MRSA, VRE |
| 126 | Actinomycin D, Actinomycin X, Antimycin, monactin | SaN, EcD, MRSA, VRE |
| 10218 | Actinomycin D, Actinomycin X | SaN, EcD, MRSA, VRE |
| 10256 | Actinomycin D, Actinomycin X | SaN, EcD, MRSA, VRE |
| 10269 | Actinomycin X | SaN, EcD, MRSA, VRE |
| 221 | Actinomycin D, Actinomycin X, antimycin | SaN, MRSA, VRE |
| 242 | Actinomycin D, Actinomycin X, Antimycin | SaN, MRSA, VRE |
| 10245 | Actinomycin D, Actinomycin X, manumycin | SaN, EcD, MRSA, VRE |
| 10356 | Actinomycin X, novobiocin | SaN, EcD, MRSA, VRE |
| 10191 | manumycin | SaN, EcD |
| 10194 | manumycin | SaN, EcD |
| 10195 | manumycin | SaN, EcD |
| 10199 | manumycin | SaN, EcD |
| 10200 | manumycin | SaN, EcD |
| 10253 | manumycin | SaN, EcD |
| 10271 | manumycin | SaN, EcD |
| 10248 | manumycin | SaN, EcD |
| 10192 | manumycin | SaN |
| 10193 | manumycin | SaN, EcD |
| 10247 | manumycin | SaN, EcD |
| 10251 | manumycin | SaN |
| 10252 | manumycin | SaN |
| 313 | monactin | SaN, EcD |
| 10174 | monactin | SaN, EcD |
| 202 | montanastatin, valinomycin | SaN, EcD, MRSA, VRE, Kp, Ab |
| 10125 | sevadicin | SaN, EcD |
| 143 | valinomycin | EcD, Ab, Kp |
| 10362 | Elaiophylin, geldanamycin, BD-12 | SaN, EcD, MRSA, VRE, EcB |
| 326 | Antimycin | SaN, EcD, MRSA, VRE |
| 303 | Antimycin | SaN |
| 105 | Antimycin | SaN, MRSA |
| 107 | Antimycin | SaN, MRSA |
| 109 | Antimycin | SaN, MRSA |

| | | |
|---|---|---|
| 118 | Antimycin | SaN, MRSA |
| 144 | Antimycin | SaN |
| 146 | Antimycin | SaN, MRSA |
| 305 | Antimycin | SaN |
| 218 | Bonactin, monactin | SaN, VRE |
| 103 | Bonactin, monactin, montanastatin, valinomycin | SaN, MRSA, VRE |
| 136 | Bonactin, monactin, montanastatin, valinomycin | SaN, MRSA, VRE |
| 201 | Bonactin, monactin, montanastatin, valinomycin | SaN, MRSA, VRE |
| 212 | Bonactin, monactin, montanastatin, valinomycin | SaN, MRSA, VRE |
| 215 | Bonactin, monactin, montanastatin, valinomycin | SaN, MRSA, VRE |
| 122 | echinomycin, tambromycin A and B | SaN, MRSA, VRE |
| 111 | monactin | SaN, MRSA, VRE |
| 125 | monactin | SaN, MRSA, VRE |
| 135 | monactin | SaN, MRSA, VRE |
| 137 | montanastatin, valinomycin | SaN, MRSA |
| 10348 | novobiocin | SaN, EcD, VRE |
| 209 | pyrromycin | SaN, MRSA, VRE |
| 210 | pyrromycin | SaN, MRSA, VRE |
| 213 | Pyrromycin | SaN, MRSA, VRE |
| 245 | Pyrromycin | SaN, MRSA, VRE |
| 203 | pyrromycin, valinomycin | SaN, MRSA, VRE |
| 217 | pyrromycin, valinomycin | SaN, MRSA, VRE |
| 133 | tambromycin A and B | SaN, MRSA |
| 106 | tambromycin A and B | SaN, VRE |
| 114 | tambromycin A and B | SaN, VRE |
| 222 | tambromycin B | SaN |
| 104 | valinomycin | SaN, MRSA |
| 204 | valinomycin | SaN, MRSA |
| 216 | valinomycin | SaN, MRSA, VRE |
| 10285 | venturicidin | SaN |

* If a strain is listed in this column, it means there was a discernible zone of clearing against that particular test isolate. Sa = *S. aureus* Newman; EcD = *E. coli* DH10B; MRSA = *S. aureus* JKD6008; VRE = *E. faecium* AUS0233; Kp = *K. pneumoniae* NTUH-K2044; Ab = *A. baumannii* BAA-1710; EcB = *E. coli* BPH0530. Refer to Table S1 for further information about each test organism.

**Table S3**. Statistics of sequenced bacterial genomes.

| Strain # | Closest 16S match (% query coverage, % nt ID) | Total length | # contigs | Largest contig | Contig N50 | # predicted SMBGC regions* |
|---|---|---|---|---|---|---|
| 10260 | *Streptomyces murinus* NBRC 100773 (97, 100%) | 8,508,414 | 76 | 569,700 | 357,808 | 36 |
| M126 | *Streptomyces antibioticus* CSSP528 (98, 100) | 9,075,813 | 196 | 360,003 | 97,734 | 32 |
| 10202 | *Streptomyces murinus* NBRC 100773 (97, 100%) | 8,454,924 | 163 | 386,948 | 166,043 | 40 |
| 10256 | *Streptomyces murinus* NBRC 100773 (97, 100%) | 8,718,858 | 2 | 8,395,928 | 8,395,928 | 45 |
| 10362 | *Streptomyces antimycoticus* NBRC 100767 (97, 99.93) | 11,796,174 | 32 | 2,605,409 | 1,130,587 | 52 |
| 10258 | *Streptomyces ardesiacus* (97, 99.26) | 8,668,535 | 4 | 8,482,255 | 8,482,255 | 29 |

* Indicates number of genomic regions with likely secondary metabolite BGCs (SMBGCs) as identified by AntiSMASH v7 (8). Due to fragmentation of some genome assemblies and the possibility that genes that form part of a single BGC may be present on multiple contigs, the number of SMBGC may be artificially inflated for some genomes. The smaller the number of contigs, the more likely this number is to be an accurate reflection of secondary metabolic capacity of an individual organism. Genomes sequences can be accessed at NCBI via BioProject PRJNA1272227.

**Table S4.** Homology between proteins encoded in actinomycin BGCs.

| Actinomycin BGC gene name | NCBI accession | Function | Orthologue locus tag in 10260 | % aa ID of 10260 orthologue* | Orthologue locus tag in M126 | % aa ID of M126 orthologue* | Orthologue in 10202 (% aa ID to ref prot) | % aa ID of 10202 orthologue* | Orthologue in 10256 | % aa ID of 10256 orthologue* |
|---|---|---|---|---|---|---|---|---|---|---|
| *acmrC* | ADG27345.1 | ATP-binding transporter | 10260_05753 | 86 | M126_03691 | 87 | 10202_00833 | 86 | 10256_00813 | 86 |
| *acmrB* | ADG27346.1 | ABC type 2 transporter | 10260_05754 | 89 | M126_03690 | 88 | 10202_00834 | 89 | 10256_00814 | 89 |
| *acmrA* | ADG27347.1 | ATP-binding transporter | 10260_05755 | 83 | M126_03689 | 86 | 10202_00835 | 83 | 10256_00815 | 83 |
| *acmQ* | ADG27348.1 | ViuB-like protein | 10260_05756 | 77 | M126_03688 | 82 | 10202_00836 | 77 | 10256_00816 | 77 |
| *acmP* | ADG27349.1 | TetR family transcriptional regulator | 10260_05757 | 71 | M126_03687 | 78 | 10202_00837 | 72 | 10256_00817 | 72 |
| *acmO* | ADG27350.1 | LbmU-like protein | 10260_05758 | 7 | M126_03686 | 76 | 10202_00838 | 70 | 10256_00818 | 70 |
| *acmN* | ADG27351.1 | ferredoxin | 10260_05760 | 88 | M126_03683 | 88 | 10202_00844 | 88 | 10256_00826 | 88 |
| *acmM* | ADG27352.1 | cytochrome P450 monooxygenase | 10260_05761 | 59 | M126_03682 | 60 | 10202_00845 | 60 | 10256_00827 | 59 |
| *acmL* | ADG27353.1 | methyltransferase | 10260_05764 | 45 | M126_03672 | 74 | 10202_00848 | 45 | 10256_00830 | 45 |
| *acmK* | ADG27353.1 | aminotransferase class V | 10260_05771 | 73 | M126_03679 | 45 | 10202_00855 | 73 | 10256_00837 | 73 |
| *acmT* | ADG27354.1 | hypothetical protein | 10260_05770 | 74 | M126_03673 | 72 | 10202_00854 | 74 | 10256_00836 | 74 |
| *acmS* | ADG27355.1 | hypothetical protein | 10260_05769 | 81 | M126_03674 | 80 | 10202_00853 | 81 | 10256_00835 | 83 |
| *acmR* | ADG27356.1 | mbtH-like protein | 10260_05768 | 72 | M126_03675 | 72 | 10202_00852 | 73 | 10256_00834 | 72 |
| *acmD* | ADG27357.1 | 4-MHA carrier protein | 10260_05767 | 72 | M126_03676 | 73 | 10202_00851 | 72 | 10256_00833 | 73 |
| *acmA* | ADG27358.1 | AMP-dependent synthetase and ligase | 10260_05766 | 73 | M126_03677 | 72 | 10202_00850 | 73 | 10256_00832 | 73 |
| *acmB* | ADG27359.1 | non-ribosomal peptide synthetase | 10260_05765 | 76 | M126_03678 | 76 | 10202_00849 | 76 | 10256_00831 | 75 |
| *acmC* | ADG27360.1 | non-ribosomal peptide synthetase | 10260_05764 | 71 | M126_03672 | 44 | 10202_00848 | 71 | 10256_00830 | 71 |
| *acmE* | ADG27360.1 | hypothetical protein | 10260_05771 | 45 | M126_03679 | 71 | 10202_00855 | 44 | 10256_00837 | 45 |
| *acmF* | ADG27361.1 | aryl formamidase | 10260_05763 | 75 | M126_03680 | 77 | 10202_00847 | 74 | 10256_00829 | 75 |
| *acmG* | ADG27362.1 | tryptophan 2,3-dioxygenase | 10260_05762 | 74 | M126_03681 | 75 | 10202_00846 | 74 | 10256_00828 | 74 |
| *acmH* | ADG27363.1 | aminotransferase class V | 10260_05761 | 84 | M126_03682 | 82 | 10202_00845 | 85 | 10256_00827 | 84 |
| *acmI* | ADG27364.1 | methyltransferase | 10260_05760 | 8 | M126_03683 | 82 | 10202_00844 | 80 | 10256_00826 | 81 |
| *acmJ* | ADG27365.1 | LbmU-like protein | 10260_05758 | 66 | M126_03686 | 71 | 10202_00838 | 66 | 10256_00818 | 66 |
| *acmU* | ADG27366.1 | TetR family transcriptional regulator | 10260_05757 | 65 | M126_03687 | 65 | 10202_00837 | 65 | 10256_00817 | 65 |

| | | | | | | | | | | |
|---|---|---|---|---|---|---|---|---|---|---|
| *acmV* | ADG27367.1 | ViuB-like protein | 10260_05756 | 76 | M126_03688 | 77 | 10202_00836 | 76 | 10256_00816 | 76 |
| *acmW* | ADG27368.1 | ATP-binding transporter | 10260_05755 | 83 | M126_03689 | 82 | 10202_00835 | 83 | 10256_00815 | 83 |
| *acmX* | ADG27369.1 | ABC type 2 transporter | 10260_05754 | 84 | M126_03690 | 85 | 10202_00834 | 84 | 10256_00814 | 84 |

* Percentage amino acid identity of the orthologue to the *act* BGC reference protein.

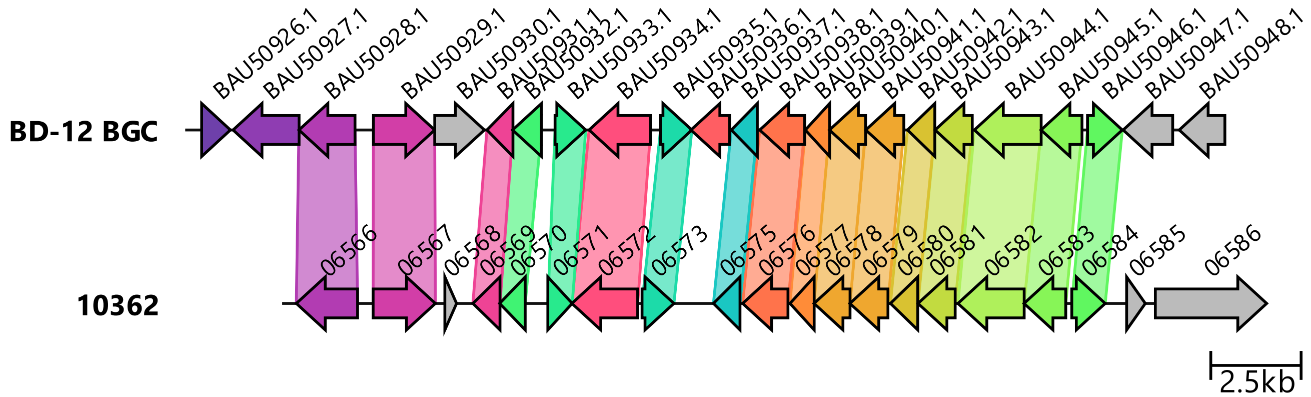


**Fig S1**. Homology between the reference BD-12 BGC (MiBIG accession BGC0001379) (9) and BD-12 BGC from strain 10362.

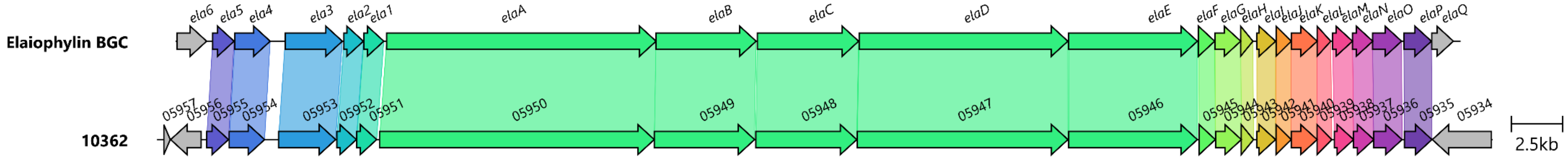


**Fig S2**. Homology between the reference elaiophylin BGC (MiBIG accession BGC BGC0000053) (10) and elaiophylin BGC from strain 10362.

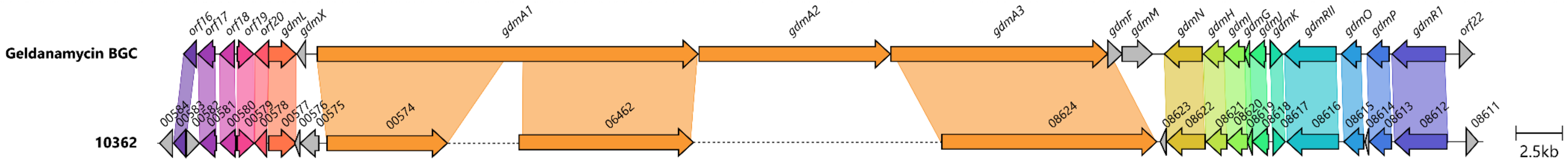


**Fig S3**. Homology between the reference geldanamycin BGC (MiBIG accession BGC0000066)(11) and geldanamycin BGC from strain 10362. The dotted line in the 10362 BGC indicates a contig break.
